# Isolation and Gene Flow in a Speciation Continuum in Newts

**DOI:** 10.1101/095877

**Authors:** Maciej Pabijan, Piotr Zieliński, Katarzyna Dudek, Michał Stuglik, Wiesław Babik

**Author notes:** Maciej Pabijan, Department of Comparative Anatomy, Institute of Zoology and Biomedical Research, Jagiellonian University, ul. Gronostajowa 9, 30-387. Wiesław Babik, Institute of Environmental Sciences, Jagiellonian University, ul. Gronostajowa 7, 30-387, Kraków, Poland.

## Abstract

Because reproductive isolation often evolves gradually, differentiating lineages may retain the potential for genetic exchange for prolonged periods, providing an opportunity to quantify and understand the fundamental role of gene flow during speciation. Here we delimit taxa, reconstruct the phylogeny and infer gene flow in newts of the *Lissotriton vulgaris* species complex based on 74 nuclear markers sampled from 127 localities. We demonstrate that distinct lineages along the speciation continuum in newts exchange nontrivial amounts of genes, affecting their evolutionary trajectories. By integrating a wide array of methods, we delimit nine taxa and show that two principle factors have driven their genetic differentiation: time since the last common ancestor determining levels of shared ancestral polymorphism, and shifts in geographic distributions determining the extent of secondary contact. Post-divergence gene flow, indicative of evolutionary non-independence, has been most extensive between sister and non-sister taxa in Central Europe, while four southern European lineages have acquired the population genetic hallmarks of independent species (*L. graecus*, *L. kosswigi*, *L. lantzi*, *L. schmidtleri*). We obtained strong statistical support for widespread mtDNA introgression, previously suggested by discordance between mtDNA phylogeny and morphology. Our study suggests that long-term evolution in structured populations that may periodically exchange genes may be common: although some of these populations may become extinct or fuse, others will acquire complete reproductive isolation and will carry signatures of this complex history in their genomes.

## 1. Introduction

Speciation, regardless of the mechanisms or geographic settings in which it occurs, is typically a gradual process (Coyne and Orr, 2004). The consequences of this are manifold. First, we see various stages of speciation in nature which makes delimitation of separately evolving units challenging (Huang and Knowles, 2016; Wiens, 2007). Second, completion of speciation is by no means certain, other outcomes such as fusions or extinction are also likely (e.g. Rhymer and Simberloff, 1996; Rudman and Schluter, 2016). Gene flow may homogenize diverging gene pools (Petit and Excoffier, 2009; Slatkin, 1985), thereby limiting or even reversing differentiation. Third, it is unclear how much gene flow occurs between differentiating lineages, whether contact between such lineages causes massive introgression, and to what extent introgression is heterogeneous throughout the genome (Cruikshank and Hahn, 2014; Wolf and Ellegren, 2016).

Although theory provides some guidelines, the preceding questions are mostly empirical and need to be addressed by studying natural systems. Recent spectacular adaptive radiations may be less informative and more exceptional in this respect than commonly assumed, because extinction may not have yet had sufficient time to prune the more ephemeral products of divergent selection (Cutter and Gray, 2016). Therefore, careful analyses of taxa composed of metapopulation lineages at various stages of divergence (de Queiroz, 2007), which can be referred to as a speciation continuum, are needed. Because the necessary analyses lie at the interface of population genetics and phylogenetics, they require integration of tools from both fields. This is especially important for studies that explicitly incorporate gene flow into the equation, because population genetic approaches for inferring post-divergence gene flow are better developed than methods co-estimating phylogeny and gene flow (Sousa and Hey, 2013). Likewise, accounting for post-divergence gene flow is a major challenge for current species delimitation methods (Petit and Excoffier, 2009).

European newts from the *Lissotriton vulgaris* group provide an empirical example of a speciation continuum influenced by genetic exchange between taxa. The Carpathian newt, *L. montandoni* (Boulenger, 1880) and the smooth newt, *Lissotriton vulgaris* (Linnaeus, 1758), have parapatric distributions in central Europe (Fig. 1a; Macgregor et al., 1990; Rafiński and Arntzen, 1987). The former is endemic to the Carpathian and easternmost Sudetes mountains, whereas the latter has a very large range in Central and Northern Europe, with a disjunct population surrounding the Greater Caucasus mountains (Fig. 1a). At the intraspecific level, *L. montandoni* is morphologically uniform across its range, reflected by only shallow genetic substructuring (Zieliński et al., 2014). In sharp contrast, *L. vulgaris* is morphologically differentiated into at least seven subspecies (Fig. 1a; Raxworthy, 1990). The smooth newt is found in a variety of habitats across the collective ranges of the subspecies (Bour et al., 2002; Bousbouras and Ioannidis, 1997; Schmidtler and Franzen, 2004; Skorinov et al., 2008) from deciduous woodlands, farmlands and pastures in European lowlands and mountainous regions, forest-steppe and taiga in western Siberia and Kazakhstan and pockets of humid habitat in otherwise dry woodland or scrub in southern Europe and Anatolia. Despite over a century of study, the status of the *L. vulgaris* subspecies is still controversial (Dubois and Raffaëlli, 2009; Schmidtler and Franzen, 2004; Speybroeck et al., 2010; Wielstra et al., 2015). The disagreement has been exacerbated by a general lack of concordance between morphologically assessed subspecific boundaries and spatial patterns of mtDNA variation (Fig. 1a, Fig. S1; Babik et al., 2005; Nadachowska and Babik, 2009; Pabijan et al., 2015). The traits used to distinguish the smooth newt subspecies pertain almost exclusively to male secondary sexual characters, which include the presence/absence and extent of dorsal crests and tail fins, toe flaps, tail filament, dorso-lateral ridges and pigmentation patterns (Raxworthy, 1990) that develop in aquatic habitats during the breeding season. Despite divergent courtship behavior (Pecio and Rafiński, 1985) that leads to strong but incomplete prezygotic sexual isolation between *L. montandoni* and *L. vulgaris* (Michalak and Rafiński, 1999; Michalak et al., 1997), these two species hybridize in and around the Carpathian mountains forming a bimodal hybrid zone (Babik and Rafiński, 2004; Babik et al., 2003; Kotlík and Zavadil, 1999). Hybridization has led to the replacement of the original mtDNA of *L. montandoni* by introgressed mtDNA of several *L. vulgaris* lineages (Babik et al., 2005; Zieliński et al., 2013). Moreover, genetic exchange among the different subspecies of *L. vulgaris* is suggested because five of the subspecific taxa do not form monophyletic mtDNA lineages (Fig. S1; Babik et al., 2005; Nadachowska and Babik, 2009; Pabijan et al., 2015). Areas of subspecies intergradation with morphological intermediates have been reported in central and southeastern Europe (e.g. Krizmanic et al., 1997; Schmidtler and Franzen, 2004; Schmidtler and Schmidtler, 1983; Wielstra et al., 2015). These findings suggest that introgressive hybridization both between *L. vulgaris* and *L. montandoni*, and among the various subspecies of *L. vulgaris*, has the potential to blur boundaries among the different taxa.

**Figure 1.**
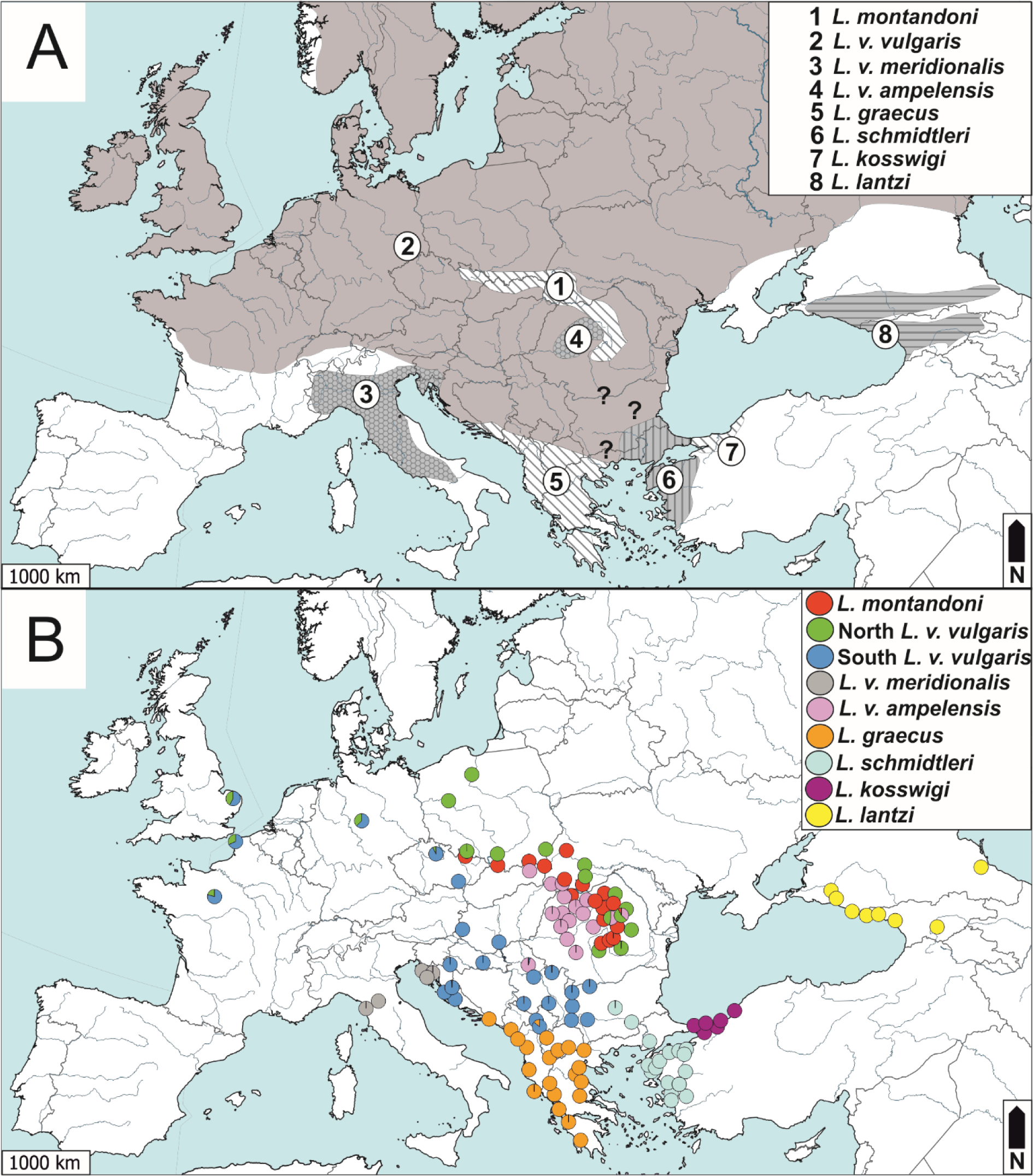
Map with ranges of morphological subspecies (A). Delimitation of evolutionary lineages in the *Lissotriton vulgaris* species group in STRUCTURE (B). Pie charts show mean individual cluster membership coefficients. *Lg – L. graecus; Lk – L. kosswigi; Ll – L. lanzti; Lm – L. montandoni; Ls – L. schmidtleri; Lva – L. v. ampelensis; Lvm – L. v. meridionalis; NLvv – North L. v. vulgaris; SLvv – South L. v. vulgaris*

Here, we provide a multi-faceted perspective on divergence and gene flow in the *L. vulgaris* complex. First, we postulate that morphologically diagnosable taxa within the complex constitute separately evolving metapopulation lineages (de Quieroz, 2007) and test this assertion using a clustering approach based on allele frequencies, a Bayesian multispecies coalescent delimitation method, and the genealogical sorting index. Second, we hypothesize that gene flow has blurred phylogenetic relationships and misled phylogeny reconstruction methods that do not take reticulation explicitly into account (Leaché et al., 2014; Solís-Lemus et al., 2016). Third, we predict that the extent of historical gene flow was most extensive among lineages inhabiting Central Europe, and use an Approximate Bayesian Computation framework to model gene flow between pairs of taxa. Finally, we apply the inferred nuclear phylogeny to statistically test the previously proposed hypothesis of extensive mtDNA introgression among morphologically diagnosable taxa. We demonstrate that isolation in space and time as well as post-divergence gene flow shape the genetic background of a set of metapopulations that have diverged in ecology and morphology including male epigamic traits.

## 2. Material and Methods

### 2.1 Newt sampling

Tissues (tail-tips) were sampled from 127 newt populations (i.e. breeding sites, Table S1 and Fig. 1b). Our sampling focused on capturing the genetic variation present in all morphologically defined taxa within the *L. vulgaris* group and all potential Pleistocene refugia (Babik et al., 2005; Nadachowska and Babik, 2009; Pabijan et al., 2015) and was therefore mostly limited to southeastern and central Europe. Because the results of the current study support the elevation of four taxa to the species level (see Discussion), we use this updated nomenclature throughout the text. One individual was sampled per locality. Altogether we analysed sequence data for 128 individuals (plus outgroup) which included 42 reported in Zieliński et al. (2016) and 87 sequenced for this study (Table S1).

### 2.2 Laboratory procedures and datasets

DNA from ethanol preserved tissues was extracted using either a standard phenol/chloroform technique or the Wizard Genomic DNA extraction kit (Promega). Amplification and sequencing of a panel of 74 nuclear, mostly 3’UTR markers, followed Zieliński et al. (2014). This procedure resulted in physically phased sequences of both haplotypes for most markers in most individuals. We used four different datasets for analyses (Table S2) i) dataset 1 with sequences of all 74 loci and all 128 individuals, ii) dataset 2 with sequences of all 74 loci but with 118 individuals (excluded 9 admixed individuals and one of two newts from locality Kapakli), iii) dataset 3 with sequences of all 128 individuals but 70 loci (excluded 4 loci with > 10% missing sequences), iv) dataset 4 consisting of SNPs extracted from 71 loci (excluded loci with >10% missing data at nucleotide positions) and 120 individuals (including outgroup).

### 2.3 Species delimitation

Species delimitation was based on identifying distinct genetic groups from haplotype frequency data and validating the genetic clusters using multispecies coalescent-based and genealogical sorting methods.

Sequence alignments (dataset 1) were converted into input for STRUCTURE v.2.3.4 (Pritchard et al., 2000) by coding each haplotype as a unique integer in a custom Python script. If nucleotide sequences contained over 10 Ns or gaps, they were coded as missing data, if they had less than 10 Ns or gaps, then columns with missing data were removed from the alignment. This procedure ensured that haplotype identity was based on nucleotide differences between sequences (and not on e.g. indels or low-quality data) and provided a conservative estimate of the number of haplotypes per locus. STRUCTURE was run under the admixture model (α = 1.0) with independent allele frequencies among populations (λ = 1.0) for *K* from 1 to 12, iterated 10 times each. The Markov chain in each analysis was set to 1 million burnin steps; a further 2 million steps were used for parameter value estimation. STRUCTURE HARVESTER (Earl and vonHoldt, 2012) and CLUMPAK (Kopelman et al., 2015) were used process the STRUCTURE output and to evaluate the most probable number of clusters (*K*) by examining logPr(*Data*).

We validated the genetic clusters identified by STRUCTURE by applying joint Bayesian species delimitation and species tree estimation using the program BPP v3.1 (Yang, 2015). This approach is based on the multispecies coalescent and compares different models of species delimitation and species phylogeny in a Bayesian framework, accounting for incomplete lineage sorting and gene tree-species tree conflict (Rannala and Yang, 2013; Yang and Rannala, 2010, 2014). Input alignments were constructed from dataset 2 by taking 10 sequences per marker for each of the 9 STRUCTURE-delimited operational taxonomic units (hereafter OTUs). This sampling strategy was based on the minimum number of sequences available per taxon (imposed by 5 specimens of *L. v. meridionalis*), with random subsampling of 10 sequences (without replacement) in all other taxa. Admixed individuals (as defined by STRUCTURE) were excluded from this analysis. We explored a wide variety of priors on the population size parameters (θs) and divergence time at the root of the species tree (τ_0_) (see Extended Methods in the Supplementary Material).

Next, we estimated gene trees in MrBayes 3.2 (Ronquist et al., 2012) and used them as input in Bayesian concordance analysis and an assessment of genealogical sorting (see below). We applied dataset 3; outgroups consisted of a single, randomly chosen haplotype per locus of a *Lissotriton helveticus* individual, or if unavailable due to lack of amplification, a single haplotype from either *L. italicus* (for 3 markers) or *L. boscai* (for 9 markers) from Zieliński et al. (2014). Nucleotide substitution models for each marker were determined in jModeltest 2.1.4 (Darriba et al., 2012) under the Bayesian information criterion. Due to the low phylogenetic information content of many of the markers, we did not partition the alignments into codon positions for those containing coding regions. We implemented two runs each with four chains for 3 million generations with trees and parameters sampled every 3000 generations. Convergence was assessed by comparing the –lnL values between runs and by examining parameter values for a subset of markers in Tracer (Rambaut and Drummond, 2007). We discarded half of recorded gene trees as burnin, combined runs and constructed an extended majority tree in MrBayes (allcompat command).

We quantified the genealogical distinctiveness of the OTUs by measuring exclusive ancestry across gene trees with the genealogical sorting index (*gsi*) of Cummings et al. (2008). This statistic measures the genealogical exclusivity of a grouping on a scale from 0 (random distribution of sequences among groups) to 1 (complete monophyly), and an ensemble statistic (*egsi*) is used to integrate across gene trees. Because the *gsi* is sensitive to disparity in group size, we randomly pruned leaves to a sample size of 10 (the minimum number of sequences available per group). Admixed individuals were also pruned from gene trees. We used the R implementation of the *gsi* (genealogicalSorting v.0.92). The null hypothesis that the degree of exclusive ancestry of each group was observed by chance was evaluated by 10^6^ permutations.

### 2.4 Phylogeny estimation

Our first phylogenetic approach involved using individuals as taxonomic units and concatenating the sequences of all loci. This procedure necessitates the selection of a single haplotype per individual per marker and becomes non-trivial if that individual is heterozygous at multiple loci, because the choice of which haplotype to concatenate into the matrix may influence both the topology and branch support of resultant trees (Weisrock et al., 2012). For each individual in dataset 2, we randomly merged haplotypes from each gene into two concatenated sequences (i.e., each individual was represented by two entries of 36, 918 bp consisting of 74 concatenated markers). We then pooled haplotypes for each candidate species and randomly selected half of the haplotypes from each group for analysis. We followed Weisrock et al. (2012) in conducting ten replicate analyses to assess consistency. We ran mixed-model phylogenetic analyses in MrBayes 3.2 on the CIPRES Science Gateway platform (Miller et al., 2010). We divided the alignments into 74 partitions corresponding to markers and assigned the optimal nucleotide substitution models to these partitions as previously determined by jModeltest. Each analysis consisted of two runs with 8 chains for 20 million generations with trees and parameters sampled every 5000 generations; burnin was set at 50%. Convergence between runs was assessed by examining effective samples sizes and parameter values between runs in Tracer, and by comparing topologies in AWTY (Nylander et al., 2008).

The second phylogenetic method involved using BUCKy (Ané et al., 2007; Larget et al., 2010) to estimate levels of concordance in reconstructed topologies among the posterior distributions of gene trees generated in MrBayes. Because the number of unique tips per gene tree was prohibitive (see Extended Methods), we pruned the gene trees in the MrBayes posterior distributions by randomly selecting a single tip from each OTU using an R script from Weisrock et al. (2012), modified to accommodate our dataset and subsampling requirements. Admixed individuals were also pruned from gene trees. We tested a range of priors and parameter values for BUCKy (see Extended Methods). We pruned the original gene trees to 10 OTUs (9 candidate species plus outgroup) and repeated the subsampling and pruning routine 50 times, using BUCKy to calculate the primary concordance trees for each replicate. We measured clade support by calculating the mean (± SD) sample-wide concordance factors for clades across all 50 replicates. We also measured clade support by computing the frequency with which each clade was recovered over all replicates.

We constructed a maximum likelihood tree using TreeMix 1.12 (Pickrell and Pritchard, 2012) which uses a covariance matrix based on allele frequencies of SNPs to infer the evolutionary relationships among pre-specified populations. The algorithm then identifies populations more closely related than predicted by the phylogeny and adds a “migration edge” between them, representing the direction and magnitude of gene flow (Pickrell and Pritchard, 2012). We used the nine OTUs and an outgroup as populations. Note that allele frequencies for our outgroup “population” were based on two gene copies because only one outgroup individual was sequenced per marker (see single gene analyses above). We extracted and sorted biallelic SNPs at each of the 74 markers according to minor allele frequency (MAF). Five SNPs with the highest MAF, i.e. most informative at the level of the entire dataset, were retrieved per marker, under the condition that they contained <10% missing data at a nucleotide position; these positions were then converted to TreeMix format using a custom Python script. A total of 71 out of 74 markers met our criteria, resulting in a matrix of 355 SNPs (dataset 4). TreeMix was invoked with –k 5 to account for linkage disequilibrium for SNPs from the same marker. In consecutive analyses, we added from one to three migration edges to allow for gene flow in the tree. We also ran TreeMix with an alternative dataset using only 1 SNP per marker with the highest MAF.

### 2.5 Gene flow among OTUs

We first summarized the pattern of shared variation in our dataset by counting the numbers of shared haplotypes between OTUs in a custom Python script using dataset 2. We also used this dataset to calculate *d*_xy_ values among the newt OTUs in DnaSP v5 (Librado and Rozas, 2009).

We estimated gene flow between all pairs of OTUs (36 pairwise comparisons) in an Approximate Bayesian Computation (ABC) framework. We followed Zieliński et al. (2016) in removing all fully coding markers from analyses, excluding all missing data by removing loci or individuals in pairwise comparisons and excluding all admixed individuals; final datasets included 60-66 loci with between 5 and 24 sequences per taxon (Table S3). The information present in the data was summarized for each pairwise comparison with a set of 9 summary statistics (Table S3 and Extended Methods).

We tested four simple scenarios of pairwise species divergence and the extent and timing of interspecific gene flow: i) a no gene flow (NGF) model in which an ancestral population splits into two at time T_SPLIT_ with no genetic exchange between descendant populations, ii) an admixture (ADM) model in which a fraction of genes is moved instantaneously from one population to the other; this can occur once independently in each direction at any time after divergence, iii) a constant gene flow (CGF) model in which gene flow is continuous and bidirectional between descendent taxa after T_SPLIT_, and iv) a recent gene flow (RGF) model in which continuous and bidirectional gene flow between descendent taxa was constrained to have occurred in the last 200,000 years (approximately 50,000 newt generations).

Coalescent simulations were done in fastsimcoal2.01 (Excoffier et al., 2013), and the ABC analysis was conducted within the ABCtoolbox (Wegmann et al., 2010); detailed prior and parameter values are given in the Extended Methods. A total of 10^6^ datasets were simulated under each demographic model from which we retained 1% (10^4^) best simulations and computed the marginal likelihood of the observed and retained datasets under the generalized linear model (Leuenberger and Wegmann, 2010). To control the type I error for multiple comparisons, we used a Bonferroni correction and removed models with the nominal P<0.0125. The best fitting model was then selected via Bayes factors.

We estimated the power to distinguish between the four models within the ABC framework by randomly picking 1000 pseudo-observed datasets from all simulations generated for each model and checking how often the ABC procedure correctly predicted the true model (the one that produced the dataset). Each pseudo-observed dataset was treated as the observed data and used to calculate marginal densities of all compared models; Bayes factors were used to select the best model. Because identical prior distributions were used for all pairwise comparisons, we conducted a single power analysis using populations with the fewest sequences (*L. kosswigi* - *L. v. meridionalis*). This can be considered the minimum estimate as all other comparisons (with larger sample sizes) will have more power in distinguishing between models.

Next, we formally tested whether the sharing of mtDNA haplotypes among *Lissotriton* groups could be accounted for by ILS (incomplete lineage sorting) or is due to hybridization. We assumed that the partially resolved nuclear phylogeny based on the concordant signal from concatenated/BUCKy/TreeMix analyses (further referred to as the species tree) reflects true relationships among the evolutionary lineages within *L. montandoni*/*L. vulgaris* and followed the statistical approach of Joyce et al. (2011). If mtDNA similarities among evolutionary lineages result from ILS, then mtDNA sequences simulated within the true species tree should replicate the observed mtDNA data. If mtDNA introgression has occurred, sequences simulated on alternative species trees, assuming closest relationships between the lineages showing highest mtDNA similarity, should better reproduce the observed data. Because the relationships between the evolutionary lineages have not been resolved completely, both tests were based on four lineages with relationships well-supported by nuclear data: i) N *L. v. vulgaris*, *L. v. ampelensis*, *L. kosswigi* and *L. montandoni*; ii) S *L. v. vulgaris*, *L. v. meridionalis*, *L. schmidtleri* and *L. graecus* (Fig. 2). For *L. montandoni* two scenarios were evaluated, allowing introgression from either *L. v. ampelensis* or N *L. v. vulgaris*. For each evolutionary lineage we randomly sampled one sequence from each locality for which mtDNA sequences were available (Babik et al., 2005; Pabijan et al., 2015; Zieliński et al., 2013). These sequences were used to calculate parameters (times of divergence and mutation-scaled effective population sizes of extant and ancestral taxa) of competing species trees in BPP v3.1. A thousand gene trees were simulated for each species tree with MCcoal. Sequences were simulated along gene trees using SeqGen (Rambaut and Grass, 1997) assuming the HKA + gamma model of sequence evolution with parameter values estimated in MEGA6 (Tamura et al., 2013). For each scenario mean and minimum uncorrected (p) distances between evolutionary lineages were calculated for 1000 simulated datasets and compared to the observed distances to obtain *P* values for individual comparisons. *P* values from individual tests were combined using Fisher’s method.

**Figure 2.**
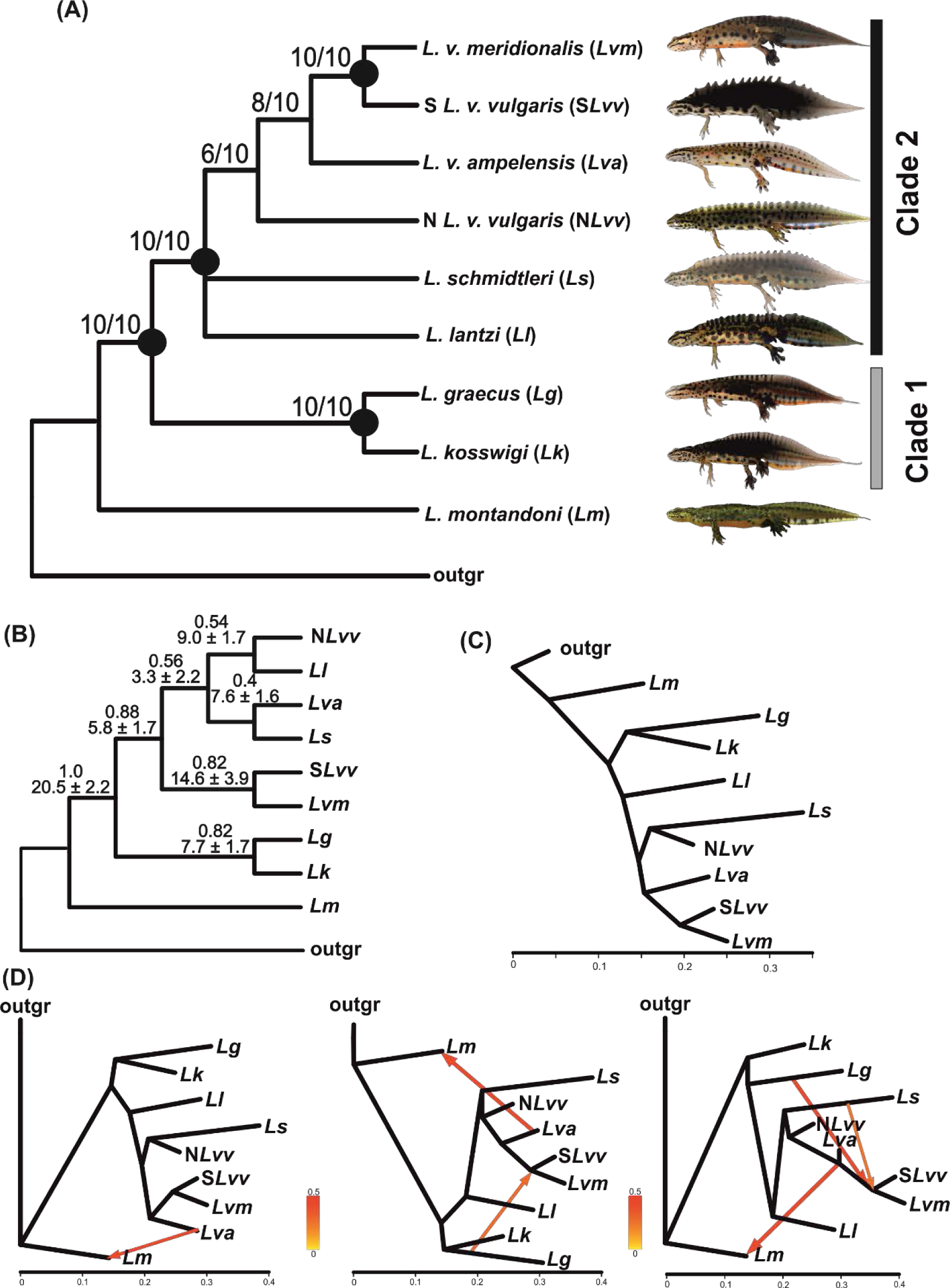
Phylogenetic relationships among STRUCTURE-delimited groups of *Lissotriton* newts; most genetic groups correspond to morphologically defined subspecies (except for *L. v. vulgaris* that is split into northeastern and southwestern lineages and *L. v. ampelensis* which contains two morphologies; see text for details). (A) Majority-rule consensus tree based on 10 replicate analyses (Fig. S4) of 74 concatenated nuclear loci for a total of 36,918 bp; values designate the frequencies of clades found across replicates. Filled circles at nodes indicate congruence across concatenation, concordance and TreeMix analyses. (B) Majority-rule consensus of 50 primary concordance trees. Frequency of clades over all replicates (top number) and concordance factors shown as the mean number of loci (out of 70) supporting the clade (± SD). (C) Maximum likelihood tree as inferred by Treemix based on 355 SNPs; horizontal branch lengths are proportional to the amount of genetic drift that has occurred in each lineage. (D) Inferred evolutionary relationships among newt groups by TreeMix with one (left), two (middle) and three (right) migration events. The migration arrows are coloured according to their weights; the weight is correlated with the ancestry fraction.

### 2.6 Data archiving

Input data and custom scripts used for data transformation and analysis are available in Dryad digital repository with doi:[to be submitted upon manuscript acceptance]. DNA sequence alignments for all markers were submitted to GenBank: accession nos. [to be submitted upon manuscript acceptance]. Additional analyses are provided as Supplementary Data.

## 3. Results

The largest dataset 1 contained 128 individuals and 74 loci with an average of 72.1 ± 20.29 haplotypes/locus (range 29-117) and 1.9 ± 3.87 % missing data per locus. Markers were of similar length (498.9 ± 6.79 bp) and highly polymorphic with an average of 91.8 ± 34.56 segregating sites per marker, one third of these (29.1 ± 9.52) were singleton polymorphisms (Table S4).

### 3.1 Species delimitation

Bayesian clustering solutions at *K* ≤ 6 had low probabilities. Pr(*K*) plateaued at *K* = 7 (Fig. S2) and therefore we further only considered analyses at *K* = 7-12 (60 analyses in total). Most of these 60 analyses contained clusters that grouped all individuals of a given subspecies or species together (Fig. 1B, Table S5). The top ten analyses with highest lnP all contained *K* = 9 groups with identical arrangements of individuals in clusters. Analyses with *K* > 9 contained at least one cluster that was assigned zero or trivial fractions of the data, indicating that under high *K* values, the number of meaningful clusters was lower than the assumed *K*. We thus inferred 9 clusters with high certainty (Fig. 1B, Table S5). Five of these clusters correspond to morphologically designated species or subspecies: *L. montandoni*, *L. graecus*, *L. v. meridionalis*, *L. kosswigi* and *L. lantzi*. The nominal subspecies was subdivided into two clusters that we designate North *L. v. vulgaris* (N *L. v. vulgaris*) and South *L. v. vulgaris* (S *L. v. vulgaris*), with the former living predominantly to the north and east of the Carpathian arc and the latter to the west and south of the Danube. The genetic cluster encompassing *L. schmidtleri*, a taxon described from western Anatolia that is morphologically similar to the nominal subspecies, was also found in the southeastern Balkans. The genetic cluster including *L. v. ampelensis* contained four individuals from eastern Slovakia, extreme western Ukraine and Romania classified as *L. v. vulgaris* according to morphology. Following Rosenberg et al. (2001), we consider that the frequency with which each taxon formed an exclusive cluster over all 60 analyses is a reliable measure of its support (Table S5). This ranged from 70% in *L. kosswigi* to 88% in *L. graecus* and *L. schmidtleri* and considerably exceeded the frequency with which particular taxa clustered together or were split into two or more groups. Some evidence of additional substructure was found in South *L. v. vulgaris* and *L. graecus* (Fig. S3).

Admixture was consistently detected in nine individuals (7.0%; Fig. 1B, see Table S1 for estimated admixture proportions), five of which originated from the western part of the range of *L. vulgaris* (admixed N *L. v. vulgaris*/S *L. v. vulgaris*, morphologically classified as *L. v. vulgaris*), a single individual from northern Serbia (*L. v. ampelensis*/S *L. v. vulgaris,* morphological *L. v. vulgaris*), one from Kosovo (S *L. v. vulgaris/L. graecus*, morphologically undetermined), and two morphologically undetermined newts from the Romanian Carpathians composed of two (N *L. v. vulgaris*/*L. v. ampelensis*) or three (N *L. v. vulgaris*/*L. v. ampelensis*/*L. montandoni*) genetic clusters.

We conducted joint inference of species delimitation and phylogeny using BPP 3 with the nine OTUs defined in STRUCTURE. The posterior probabilities of each of these equaled 1.00 in all analyses under a wide range of priors and different starting trees (Table 1). All models in the 95% credibility sets contained the nine OTUs. The multispecies coalescent model in BPP recovered a number of different species trees under the applied parameter settings (Table 1); their topologies and posterior probabilities varied depending on the prior specification, and to a limited extent, on seed number. Although the latter may suggest a lack of convergence, species trees with highest support were mostly congruent across analyses. Notably, *L. montandoni* and *L. v. ampelensis* were consistently recovered as sister taxa.

**Table 1.** Joint Bayesian species delimitation and species tree estimation in the *Lissotriton vulgaris* species group using BPP. Rows show results for different analyses; letters denote replicate runs differing only by seed number. Population size (θ) and divergence time (τ) priors encompass a wide range of demographic and divergence scenarios. Starting guide tree topologies were random unless specified otherwise (see footnotes). *N* species shows the numbers of species and their posterior probabilities (PPs) in each analysis (PPs for each species within analyses were always 1.00). *N* trees denotes the number of trees in the 95% credibility set, and the last column shows the topology and posterior probability of the highest ranked tree in the set.

| Runs | $\Theta^a$ | $\tau^a$ | *N* species (PP) | *N* trees | best tree and pp of model (delimitation + tree) |
| --- | --- | --- | --- | --- | --- |
| 1a | G(1, 35) | G(2, 500) | 9 (1.00) | 7 | (*L. graecus*, (*L. lantzi*, ((*L. schmidtleri*, *L. kosswigi*), ((*S L. v. vulgaris*, *L. v. meridionalis*), (*N L. v. vulgaris*, (*L. montandoni*, *L. v. ampelensis*)))))); 0.70905 |
| 1b | G(1, 35) | G(2, 500) | 9 (1.00) | 4 | (*L. graecus*, (*L. lantzi*, (*L. kosswigi*, ((*S L. v. vulgaris*, *L. v. meridionalis*), (*L. schmidtleri*, (*N L. v. vulgaris*, (*L. montandoni*, *L. v. ampelensis*)))))); 0.83335 |
| 2a<sup>b</sup> | G(1, 35) | G(2, 500) | 9 (1.00) | 4 | (*L. graecus*, (*L. lantzi*, (*L. kosswigi*, ((*S L. v. vulgaris*, *L. v. meridionalis*), (*L. schmidtleri*, (*N L. v. vulgaris*, (*L. montandoni*, *L. v. ampelensis*)))))); 0.81770 |
| 2b<sup>b</sup> | G(1, 35) | G(2, 500) | 9 (1.00) | 2 | (*L. graecus*, (*L. lantzi*, (*L. kosswigi*, ((*S L. v. vulgaris*, *L. v. meridionalis*), (*L. schmidtleri*, (*N L. v. vulgaris*, (*L. montandoni*, *L. v. ampelensis*)))))); 0.88070 |
| 3a<sup>c</sup> | G(1, 35) | G(2, 500) | 9 (1.00) | 3 | (*L. graecus*, (*L. lantzi*, (*L. kosswigi*, ((*S L. v. vulgaris*, *L. v. meridionalis*), (*L. schmidtleri*, (*N L. v. vulgaris*, (*L. montandoni*, *L. v. ampelensis*)))))); 0.82170 |
| 3b<sup>c</sup> | G(1, 35) | G(2, 500) | 9 (1.00) | 2 | (*L. graecus*, (*L. lantzi*, (*L. kosswigi*, ((*S L. v. vulgaris*, *L. v. meridionalis*), (*L. schmidtleri*, (*N L. v. vulgaris*, (*L. montandoni*, *L. v. ampelensis*)))))); 0.85825 |
| 4a<sup>d</sup> | G(1, 35) | G(2, 500) | 9 (1.00) | 3 | (*L. graecus*, (*L. lantzi*, (*L. kosswigi*, ((*S L. v. vulgaris*, *L. v. meridionalis*), (*L. schmidtleri*, (*N L. v. vulgaris*, (*L. montandoni*, *L. v. ampelensis*)))))); 0.79815 |
| 4b<sup>d</sup> | G(1, 35) | G(2, 500) | 9 (1.00) | 38 | (*L. graecus*, (*L. lantzi*, (*L. kosswigi*, ((*S L. v. vulgaris*, *L. v. meridionalis*), (*L. schmidtleri*, (*N L. v. vulgaris*, (*L. montandoni*, *L. v. ampelensis*)))))); 0.43710 |
| 5a | G(1, 10) | G(2, 2000) | 9 (1.00) | 9 | (*L. graecus*, (*L. lantzi*, (*L. kosswigi*, (*L. v. meridionalis*, (*L. schmidtleri*, ((*L. montandoni*, *L. v. ampelensis*), (*N L. v. vulgaris*, *S L. v. vulgaris*)))))); 0.33805 |
| 5b | G(1, 10) | G(2, 2000) | 9 (1.00) | 5 | (*L. graecus*, (*L. lantzi*, (*L. kosswigi*, ((*S L. v. vulgaris*, *L. v. meridionalis*), (*L. schmidtleri*, (*N L. v. vulgaris*, (*L. montandoni*, *L. v. ampelensis*)))))); 0.71930 |
| 6a | G(2,2000) | G(2,2000) | 9 (1.00) | 50 | (*L. v. meridionalis*, (((*L. graecus*, *L. lantzi*), (*S L. v. vulgaris*, (*L. montandoni*, *L. v. ampelensis*))), (*N L. v. vulgaris*, (*L. schmidtleri*, *L. kosswigi*))); 0.13655 |
| 6b | G(2,2000) | G(2,2000) | 9 (1.00) | 45 | ((*L. graecus*, *L. lantzi*), ((*L. schmidtleri*, *L. kosswigi*), ((*S L. v. vulgaris*, *L. v. meridionalis*), (*N L. v. vulgaris*, (*L. montandoni*, *L. v. ampelensis*))))); 0.12370 |
| 7a | G(1, 10) | G(1, 10) | 9 (1.00) | 3 | (*L. graecus*, (*L. lantzi*, (*L. kosswigi*, ((*S L. v. vulgaris*, *L. v. meridionalis*), (*L. schmidtleri*, (*N L. v. vulgaris*, (*L. montandoni*, *L. v. ampelensis*)))))); 0.66405 |
| 7b | G(1, 10) | G(1, 10) | 9 (1.00) | 5 | (*L. graecus*, (*L. lantzi*, ((*L. schmidtleri*, *L. kosswigi*), ((*S L. v. vulgaris*, *L. v. meridionalis*), (*N L. v. vulgaris*, (*L. montandoni*, *L. v. ampelensis*)))))); 0.77125 |
| 8a | G(1, 10) | G(2,1000) | 9 (1.00) | 3 | (*L. graecus*, (*L. lantzi*, ((*L. schmidtleri*, *L. kosswigi*), ((*S L. v. vulgaris*, *L. v. meridionalis*), (*N L. v. vulgaris*, (*L. montandoni*, *L. v. ampelensis*)))))); 0.80270 |
| 8b | G(1, 10) | G(2,1000) | 9 (1.00) | 9 | (*L. graecus*, (*L. lantzi*, (*L. kosswigi*, ((*S L. v. vulgaris*, *L. v. meridionalis*), (*L. schmidtleri*, (*N L. v. vulgaris*, (*L. montandoni*, *L. v. ampelensis*)))))); 0.48025 |

<sup>a</sup> - G(1, 35), G(2, 500): empirically derived priors; G(1, 10), G(2, 2000): large populations
with shallow divergence; G(2,2000), G(2,2000): small populations with shallow divergence;
G(1, 10), G(1, 10): large populations with deep divergence; G(1, 10), G(2,1000): large
populations with intermediate divergence
<sup>b</sup> - starting tree according to consensus phylogeny
<sup>c</sup> - starting tree reflects gene flow among *L. montandoni*, *L. v. ampelensis* and *N L. v. vulgaris*
<sup>d</sup> - starting tree reflects gene flow among *N L. v. vulgaris* and *S L. v. vulgaris*, basal position
of *L. lantzi*

We used the gene trees calculated in MrBayes to estimate the genealogical exclusivity of each candidate species. Typically, each genealogy in the post-burnin posteriors had a different topology. This was expected because there was limited phylogenetic information contained within each marker and many tips (257 including outgroup) in the trees. Unsurprisingly, consensus gene trees calculated for each marker were mostly unresolved, and showed high levels of discordance (data not shown). Nonetheless, the majority of the genealogies showed a nonrandom clustering pattern for the delimited *Lissotriton* groups as indicated by the highly significant *egsi* (Table 2). We inferred high exclusivity for *L. kosswigi* and *L. lantzi* and with about 40% of genes monophyletic, and moderate *egsi* values for *L. graecus*, *L. montandoni*, *L. schmidtleri* and *L. v. meridionalis*. The values of *egsi* were relatively low but still highly significant for *L. v. ampelensis*, S *L. v. vulgaris* and N *L. v. vulgaris*.

**Table 2.** Ensemble genealogical sorting index (*egsi*) for *Lissotriton* operational taxonomic units (10 alleles per group) based on 70 gene trees. All of the P values for *egsi* are <0.000001. The last column provides the number of gene trees in which a group reached complete monophyly.

| **group** | **egsi** | **monophyly** |
| --- | --- | --- |
| *L. lantzi* | 0.692 | 27 |
| *L. kosswigi* | 0.679 | 30 |
| *L. graecus* | 0.564 | 17 |
| *L. schmidtleri* | 0.559 | 14 |
| *L. montandoni* | 0.548 | 16 |
| *L. v. meridionalis* | 0.497 | 7 |
| *L. v. ampelensis* | 0.349 | 0 |
| South *L. v. vulgaris* | 0.343 | 0 |
| North *L. v. vulgaris* | 0.281 | 0 |

### 3.2 Phylogenetic results

In concatenation analyses, we found that the individuals within OTUs always formed clades with posterior probabilities (PP) of 1.0, we therefore collapsed individuals into groups and show a consensus tree over all replicates (Fig. 2A) and separately for each of the 10 concatenation replicates (Fig. S4). The positions of some clades were consistent across all replicates and were always fully supported (PP = 1.0). These included the basal placement of *L. montandoni,* the subsequent split of clade 1 composed of *L. graecus* and *L. kosswigi*, and the sister group relationship of *L. v. meridionalis* and S *L. v. vulgaris*. Moreover, the concatenated analyses invariably identified clade 2 composed of *L. lantzi*, *L. schmidtleri*, S *L. v. vulgaris*, N *L. v. vulgaris*, *L. v. ampelensis* and *L. v. meridionalis*. Our replicate analyses differed in the choice of sequences at heterozygous loci during the concatenation process; the results show that this influenced the branching patterns of the concatenated phylogenetic analyses by altering the positions of 4 taxa within clade 2 (*L. lantzi*, *L. schmidtleri*, N *L. v. vulgaris* and *L. v. ampelensis*). Nearly all alternative arrangements were strongly supported in one or more concatenation replicates (Fig. S4).

A consensus of the 50 replicate primary concordance trees from BUCKy (Fig. 2B) shows that all *L. vulgaris* taxa grouped together (to the exclusion of *L. montandoni*) across all replicates. The mean concordance factor represented by the mean number of genes supporting this clade was 20.5 ± 2.2 out of a total of 70 genes. Clades 1 and 2 as well as the sister group relationship between S *L. v. vulgaris* and *L. v. meridionalis* were relatively well-supported, although concordance factors were generally low (on average <15 genes), suggesting that more than one tree may equally well describe the history of these taxa. We examined this issue by extracting concordance factors from each replicate BUCKy analysis for clades with relatively high support (mostly encompassing those with two terminal taxa, Fig. S5). Primary histories were well-discernible in the cases of *L. montandoni*, *L. v. meridionalis*, S *L. v. vulgaris*, *L. lantzi*, *L. graecus* and *L. kosswigi*. However, between 3 and 6 primary histories could be discerned for N *L. v. vulgaris*, *L. v. ampelensis* and *L. schmidtleri*, shown by similar mean concordance factors and largely overlapping quartiles (Fig. S5).

The maximum likelihood topologies recovered by TreeMix based on 1 or 5 SNPs per marker (Fig. 2C) were very similar to the consensus topology of the concatenated analysis (Fig. 2A). After adding migration edges, we recovered gene flow between *L. v. ampelensis* and *L. montandoni*, followed by gene flow of lesser magnitude between *L. graecus* and *L. schmidtleri* and the ancestor of S *L. v. vulgaris* and *L. v. meridionalis* (Fig. 2D and Fig. S6).

### 3.3 Gene flow among lineages

We found a gradient in divergence values (*d*_xy_) among newt OTUs ranging between 1.14-1.59% (Fig. 3, Table S6). Each pair of OTUs shared haplotypes at some markers (Fig. 3), although in many cases this amounted to less than 2% and may be indicative of retained ancestral variation. The highest proportions were shared between N *L. v. vulgaris* and *L. v. ampelensis* (9.1%), *L. montandoni* and two other lineages (7.5 and 6.7% for N *L. v. vulgaris-L. montandoni* and *L. v. ampelensis*-*L. montandoni*, respectively) and three pairs involving S *L. v. vulgaris* and neighboring clades *L. v. ampelensis*, N *L. v. vulgaris* and *L. v. meridionalis* (5.2, 4.6 and 3.3%, respectively). We note that all instances of relatively high levels of haplotype sharing were inferred for Central European taxa that at present occur parapatrically.

**Figure 3.**
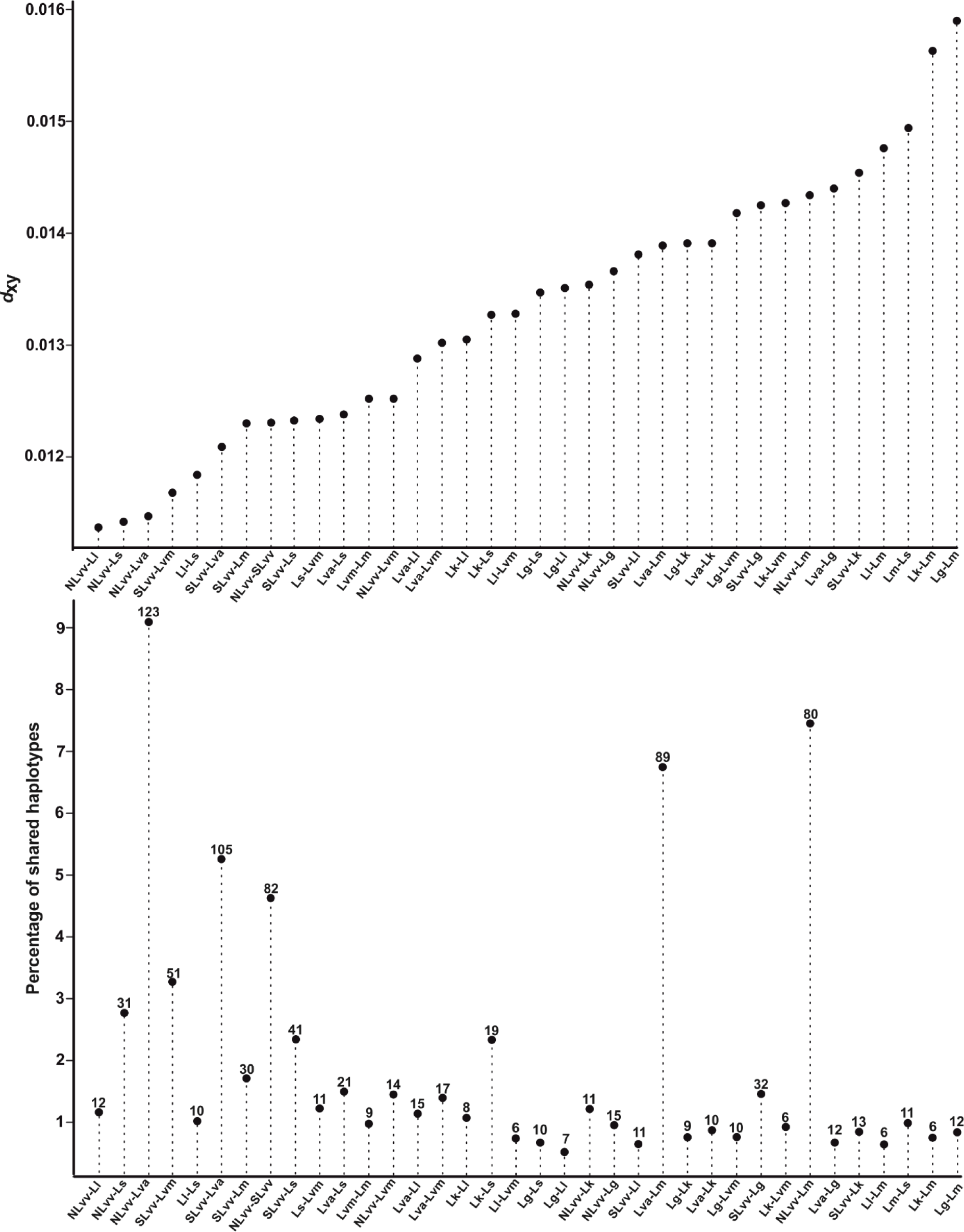
Top: pairwise *d*_xy_ distances among *Lissotriton* taxa. Bottom: proportion (in percent, left y-axis and solid circles) of shared haplotypes out of total number of haplotypes summed for each taxon pair across 74 markers; values above solid circles indicate the number of alleles shared by each taxon pair. Note that taxon pairs are in the same order on both charts. *Lg – L. graecus; Lk – L. kosswigi; Ll – L. lanzti; Lm – L. montandoni; Ls – L. schmidtleri; Lva – L. v. ampelensis; Lvm – L. v. meridionalis; NLvv – North L. v. vulgaris; SLvv – South L. v. vulgaris*

In most pairwise comparisons of OTUs within the ABC framework at least one model could reproduce the observed data (Tables S7 and S8). The analyses had good power to discriminate between the four models (Table S8). Posterior probabilities for models varied considerably among pairwise taxa comparisons (Fig. 4A) with a clear predominance of the admixture (ADM) model and only 6 comparisons in which recent (in the last 200 kya) or continuous gene flow prevailed, all confined to currently parapatric, Central European lineages. However, in other cases episodes of gene flow in the ADM model were timed to have occurred deep in the past (e.g. modal values for posteriors >1 Ma), making ADM and NGF equivalent in the sense of ruling out recent and constant gene flow (e.g. in *L. kosswigi*-*L. v. meridionalis*, *L. kosswigi*-*L. graecus, L. kosswigi-L. lantzi, L. montandoni-L. lantzi, L. montandoni-*S *L. v. vulgaris*; Table S9). In other pairwise comparisons modal values for admixture times in the ADM model were inferred to have occurred relatively recently, close to the upper limit of the prior (e.g. *L. montandoni*-*L. v. ampelensis*, N *L. v. vulgaris*-*L. kosswigi*), making the ADM and RGF models comparable. However, the posterior distributions for the admixture time parameters were in general very broad, indicating that data were not very informative in dating admixture events.

**Figure 4.**
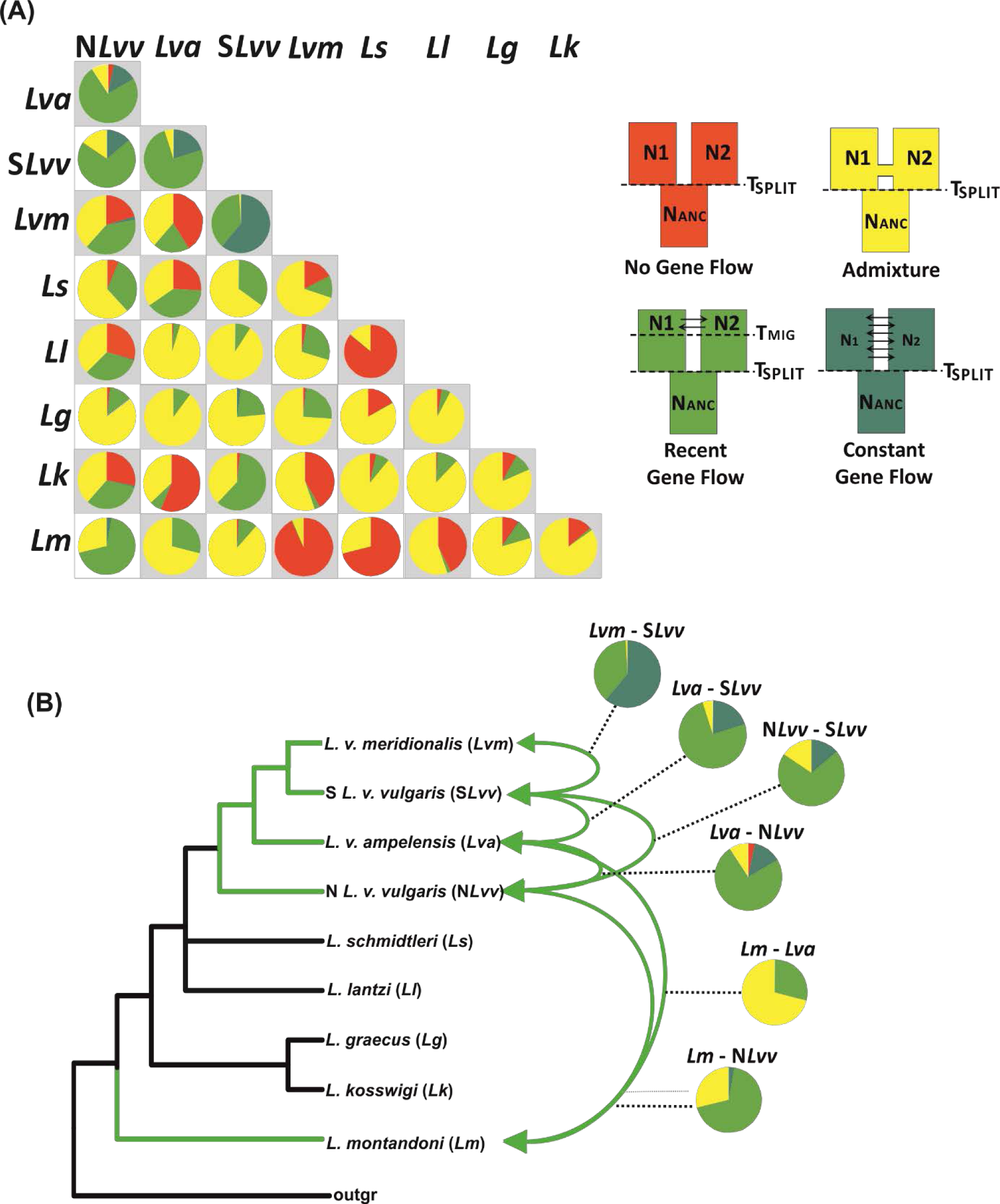
Pie charts representing posterior distributions of Approximate Bayesian Computation analyses in pairwise taxon comparisons (A), color coded according to demographic models (inset in A). *Lissotriton vulgaris* group phylogeny showing well-supported instances of post-divergence gene flow (green arrows) and their corresponding posterior distributions from the ABC analyses (B).

Phylogenies estimated from mtDNA under the assumption of ILS substantially differed from trees estimated assuming mtDNA introgression (i.e. the topology was incongruent with the species tree based on nuclear data, Fig. 5). Under ILS entire trees and internal branches were both very short. Minimum mtDNA divergence between lineages readily distinguished between ILS and hybridization. For both *L. montandoni* and *L. graecus*, scenarios assuming hybridization fit the data well and were strongly preferred over ILS (Table 3). One of the two hybridization scenarios involving *L. montandoni* fit the data better than the other, although both hybridization scenarios had much better fit than ILS. Apparently the scenario assuming introgression between N *L. v. vulgaris* and *L. montandoni* received higher support because of more extensive mtDNA haplotype sharing between these two than between *L. montandoni* and *L. v. ampelensis*. Mean mtDNA divergence between lineages was not informative in distinguishing between ILS and hybridization as both scenarios generated mean divergence that was not significantly different from the observed values (data not shown).

**Figure 5.**
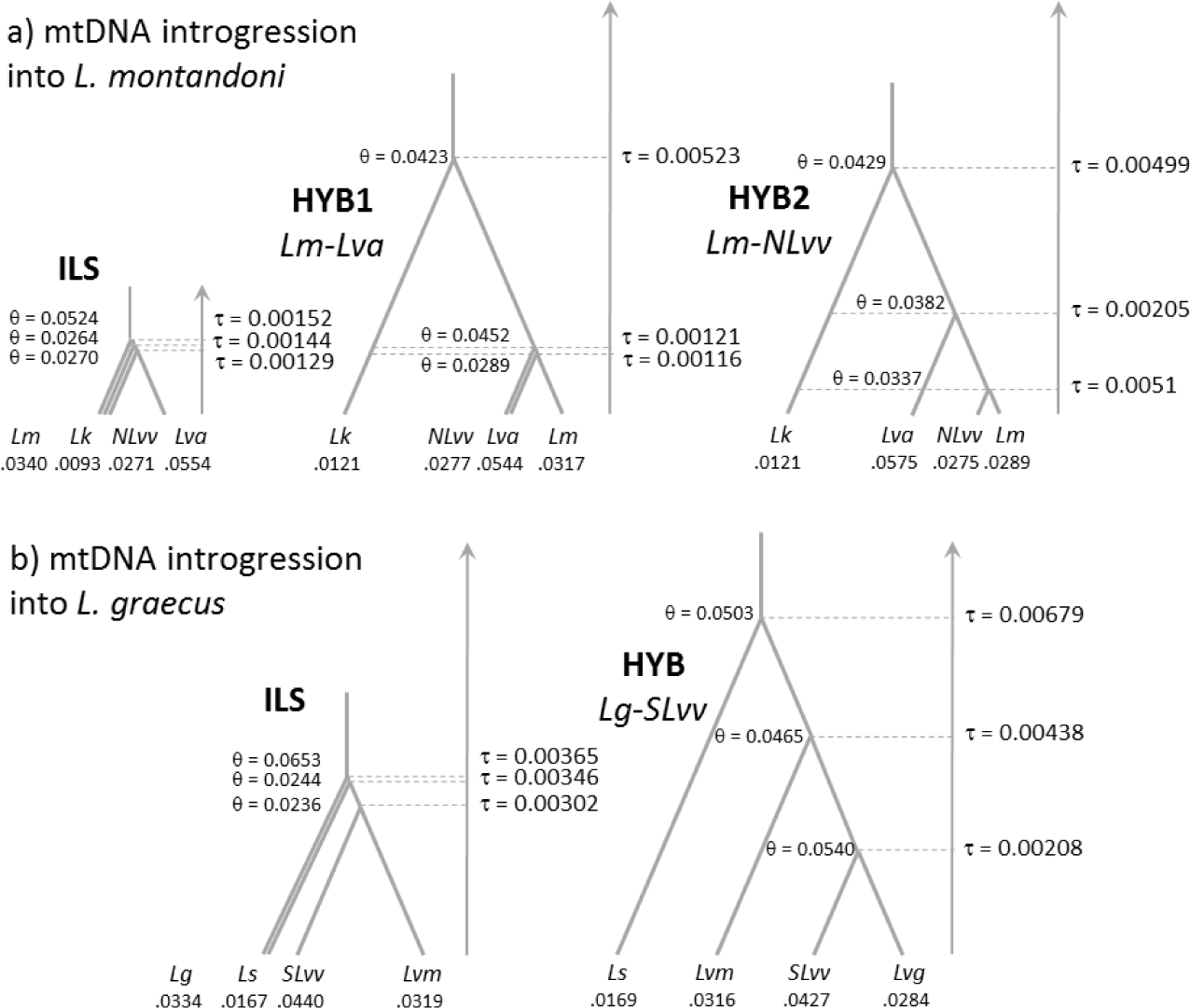
Parameters of species trees under competing scenarios of incomplete lineage sorting (ILS) and hybridization (HYB). Both θ and τ are measured as the expected number of mutations per site.

**Table 3.** Evaluation of fit of mtDNA sequence data to scenarios of incomplete lineage sorting (ILS) and hybridization (Hyb). *Lva* – *L. v. ampelensis*; *Lg* – *L. graecus*; *Lk* – *L. kosswigi; Lvm* – *L. v. meridionalis; Lm* – *L. montandoni*; *Ls* – *L. schmidtleri*; N *Lvv* – North *L. v. vulgaris*; S *Lvv* – South *L. v. vulgaris*

| Scenario | Pair of lineages | Minimum divergence - Observed | Minimum divergence - Expected (median of 1000 simulations) | P | Interpretation |
| --- | --- | --- | --- | --- | --- |
| **mtDNA introgression into *L. montandoni*** |  |  |  |  |  |
| ILS | *Lm-Lva* | 0.0020 | 0.0010 | 0.258 | O = E |
|  | *Lm -NLvv* | 0.0000 | 0.0020 | 0.294 | O = E |
|  | *Lm -Lk* | 0.0384 | 0.0020 | 0.002 | O > E |
|  | *Lva-NLvv* | 0.0020 | 0.0010 | 0.220 | O = E |
|  | *Lva-Lk* | 0.0207 | 0.0020 | 0.006 | O > E |
|  | *NLvv-Lk* | 0.0374 | 0.0030 | 0.002 | O > E |
|  | Combined (Fisher's method) |  |  | 0.00002 | data do not fit scenario |
| Hyb1<br>*Lm -Lva* | *Lm -Lva* | 0.0020 | 0.0010 | 0.034 |  |
|  | *Lm -NLvv* | 0.0000 | 0.0010 | 0.500 | O = E |
|  | *Lm -Lk* | 0.0384 | 0.0098 | 0.046 | O > E |
|  | *Lva-NLvv* | 0.0020 | 0.0010 | 0.242 | O = E |
|  | *Lva-Lk* | 0.0207 | 0.0098 | 0.150 | O = E |
|  | *NLvv-Lk* | 0.0374 | 0.0108 | 0.066 | O = E |
|  | Combined (Fisher's method) |  |  | 0.0095 | data do not fit scenario |
| Hyb2<br>*Lm -NLvv* | *Lm -Lva* | 0.0020 | 0.0020 | 0.514 | O = E |
|  | *Lm -NLvv* | 0.0000 | 0.0000 | 0.318 | O = E |
|  | *Lm -Lk* | 0.0384 | 0.0089 | 0.040 | O > E |
|  | *Lva-NLvv* | 0.0020 | 0.0020 | 0.824 | O = E |
|  | *Lva-Lk* | 0.0207 | 0.0089 | 0.110 | O = E |
|  | *NLvv-Lk* | 0.0374 | 0.0098 | 0.056 | O = E |
|  | Combined (Fisher's method) |  |  | 0.0561 | data fit scenario |
| **mtDNA introgression into *L. graecus*** |  |  |  |  |  |
| ILS | *Lg -SLvv* | 0.0000 | 0.0049 | 0.006 | O < E |
|  | *Lg -Lvm* | 0.0139 | 0.0059 | 0.120 | O = E |
|  | *Lg -Ls* | 0.0208 | 0.0059 | 0.036 | O > E |
|  | *SLvv-Lvm* | 0.0069 | 0.0039 | 0.164 | O = E |
|  | *SLvv-Ls* | 0.0139 | 0.0049 | 0.042 | O > E |
|  | *Lvm-Ls* | 0.0396 | 0.0069 | 0.022 | O > E |
|  | Combined (Fisher's method) |  |  | 0.0001 | data do not fit scenario |
| Hyb<br>*Lg-SLvv* | *Lg-SLvv* | 0.0000 | 0.0020 | 0.180 | O = E |
|  | *Lg-Lvm* | 0.0139 | 0.0069 | 0.086 | O = E |
|  | *Lg-Ls* | 0.0208 | 0.0118 | 0.212 | O = E |
|  | *SLvv-Lvm* | 0.0069 | 0.0069 | 0.832 | O = E |
|  | *SLvv-Ls* | 0.0139 | 0.0118 | 0.560 | O = E |
|  | *Lvm-Ls* | 0.0396 | 0.0138 | 0.080 | O = E |
|  | Combined (Fisher's method) |  |  | 0.1152 | data fit scenario |

## 4. Discussion

This study documents a series of lineages representing a continuum of genetic divergence with ample evidence for historical contact and gene flow between lineages. We demonstrate the importance of geographic conditions (allopatry vs. parapatry) and historical contingencies such as time since divergence and timing of secondary contact in determining the continuum of divergence that can be quantified in the present. Moreover, we show that a combination of population genetic and phylogenetic approaches is required to describe and measure this continuum. In the following sections we discuss our findings for *Lissotriton* newts, their more general implications and outline research areas where methodological advances are needed.

## 4.1 Robust delimitation despite substantial gene flow

All applied delimitation methods, based on allele frequencies, multispecies coalescent and genealogical concordance, robustly delimited nine evolutionary lineages. Some of these lineages correspond to described morphological subspecies, while others are morphologically cryptic (N *L. v. vulgaris*, S *L. v. vulgaris*) or include two distinct morphologies (*L. v. ampelensis*). This result is not an artifact of limited sampling under isolation by distance, because admixed individuals are rare and boundaries between lineages in well-sampled areas are sharp. The delimited entities are distinguishable in both allele frequency-based and genealogical frameworks, but are at various stages of divergence (Fig. 3). Substantial admixture between N *L. v. vulgaris* and S *L. v. vulgaris* in the postglacial expansion area in western Europe indicates that some lineages retain considerable potential for genetic exchange. We found evidence for recent gene flow between five other lineage pairs, all of which occur parapatrically in Central Europe (Figs 1 and 4b; see Zieliński et al., 2016, for a detailed assessment of genetic exchange between *L. montandoni* and two *L. vulgaris* lineages). Moreover, episodes of gene flow between lineages in the more distant past may have been commonplace, given the overall weak support for the divergence with no gene flow model in the ABC analyses (Fig. 4), two cases of confirmed old mtDNA introgression (Table 3), and several other instances of mtDNA paraphyly (Babik et al., 2005; Nadachowska and Babik, 2009; Pabijan et al., 2015). Thus, it is by no means certain that all delimited newt lineages will continue to evolve independently. The newt system is a good example of a broader phenomenon: as reproductive isolation progresses, the incipient species experience a prolonged phase of parapatric/allopatric structured populations which may episodically exchange genes. In some instances, secondary contact may lead to the fusion of non-sister lineages over parts of their collective range (e.g. N *L. v. vulgaris* and S *L. v. vulgaris* in western Europe). This interpretation is in line with recent genetic and molecular studies of speciation showing that post-divergence gene flow is common, e.g. it was an important feature of hominin history (Pääbo, 2015), and sometimes occurs at high levels (e.g. Muir et al., 2012; Nevado et al., 2011; Osborne et al., 2016; Sun et al., 2012).

## 4.2 Newt phylogeny and evolution of male nuptial morphology

The species tree of the *L. vulgaris* group could not be fully resolved (Fig. 2A). Relationships that received high support included *L. montandoni* as the outgroup to all *L. vulgaris* taxa, *L. graecus* and *L. kosswigi* (clade 1) as the sister group to other *L. vulgaris* lineages (clade 2), and the sister group relationship between S *L. v. vulgaris* + *L. v. meridionalis*. Although the monophyly of clade 2 is supported, the relationships among OTUs within this group are uncertain. However, we acknowledge that only one (TreeMix) of the four applied phylogenetic methods specifically modelled gene flow among OTUs. Nonetheless, well-supported nodes in the concatenation and concordance analyses were compatible with the topology of the maximum likelihood tree from TreeMix, suggesting that gene flow among OTUs did not overwhelm the phylogenetic signal over all parts of the tree. However, species tree estimation in BPP under the multispecies coalescent consistently recovered a sister group relationship between *L. montandoni* and *L. v. ampelensis* (Table 1). Likewise, analyses using *BEAST (Heled and Drummond, 2010), which experienced overall convergence issues, produced similar results (data not shown). Given the strong evidence for recent gene flow among these two taxa (Fig. 4; Zieliński et al., 2016), we consider this relationship as highly unlikely. Coalescent-based species tree methods are sensitive to violations of the assumption of no gene flow, particularly between non-sister taxa (Leaché et al., 2013; Solís-Lemus and Ané, 2016), and are thus unsuited to study the relationships within the *L. vulgaris* complex. Given the increasing number of studies reporting reticulate evolution, there is an urgent need for progress in methods co-estimating phylogeny and gene flow in multispecies and multilocus datasets; phylogenetic multilocus network methods (Solís-Lemus and Ané, 2016; Yu et al., 2013, 2014) appear promising in this respect.

The species tree for the *L. vulgaris* group nonetheless sheds light on morphological evolution in newts. The male nuptial morphology of *L. montandoni* consists of a smooth and low crest, prominent tail filament and dorso-lateral ridges. Similar male morphologies are also present in the more distantly related *L. helveticus*, *L. italicus* and *L. boscai*. This particular suite of phenotypic traits may be an adaptation for the effective transmission of waterborne pheromones (Pecio and Rafiński, 1985) and could signify the predominance of olfactory cues for species recognition (Secondi et al., 2005). On the other hand, the derived condition of prominent crests and overall larger male body size in most *L. vulgaris* lineages indicates a shift towards visual cues (Secondi et al., 2012), although other explanations are possible such as divergence in female preference for larger male body size (Haerty et al., 2007; Secondi et al., 2010). Whatever the underlying cause(s), our species phylogeny shows that *montandoni-*like male morphology was present in the ancestral *L. vulgaris* population, conserved in the *L. graecus*/*L. kosswigi* clade, and replaced by the derived morphology in the sister clade giving rise to other extant *L. vulgaris* lineages. The intermediate breeding male morphologies of *L. v. ampelensis* and *L. v. meridionalis* (Raxworthy, 1990) could be interpreted as independent, partial reversions to the ancestral morphology due to convergent evolution. Alternatively, and possibly more likely given the evidence for substantial gene flow from neighboring taxa, these intermediate morphologies could be the result of an influx of genes underlying the derived morphology and dilution of *montandoni*-like ancestral traits. These two hypotheses could be tested with genome-wide data.

## 4.3 Isolation and gene flow revealed by ABC modeling

Recognizing the limitations of existing methodologies, we explored gene flow between pairs of taxa in a model-based framework using Approximate Bayesian Computation (ABC). This approach, although quite simple, has been effective in unraveling the process of gene flow in the newt system. The data had good power in distinguishing among the four scenarios of gene flow and provided quantitative measures of support for the various models. However, analyses modeling gene flow between two populations without incorporation of additional populations have been criticized because ignoring such “ghost” populations may affect parameter estimates or even model choice (Beerli, 2004; Eaton et al., 2015; Slatkin, 2005; Strasburg and Rieseberg, 2010). We note though that pairwise comparisons among all extant taxa may partially alleviate this issue. The availability of all pairwise comparisons allows the identification and reinterpretation of problematic cases, such as support for recent gene flow between allopatric taxa which have not likely been in recent contact, but which both may have exchanged genes with other populations. In our case, inferences were largely consistent with the geographic distributions of the lineages (parapatry vs. allopatry) and we accept them as working hypotheses. In general, given the current state of the field, the use of simple pairwise models in complexes of interbreeding species may be useful because they offer a feasible way of distinguishing among different scenarios of gene flow and building more complex, but still testable, hypotheses.

In addition to model selection, ABC estimates demographic parameters which facilitate the interpretation of evolutionary history along the speciation continuum. Estimates of long-term, coalescent effective population sizes (N_e_) for the *Lissotriton* lineages are quite large, ranging roughly between 80,000 and 800,000 (Table S9). Independent assessments place the divergence time of *L. montandoni* and *L. vulgaris* between 3.7 to 6.3 mya (Stuglik and Babik, 2016), while mtDNA lineages within *L. vulgaris* split in close succession during the late Pliocene and early Pleistocene (Babik et al., 2005; Pabijan et al., 2015). This timeframe suggests that most *Lissotriton* lineages have not attained the 4N_e_ generations when roughly half of the loci in the genome achieve reciprocal monophyly (Hudson and Coyne, 2002; Degnan and Rosenberg, 2009), thus widespread incomplete lineage sorting (Table 2) and the presence of identical haplotypes amongst all lineage pairs are expected (Fig. 3). However, the lineages exchanging genes most extensively (*L. v. ampelensis*, N *L. v. vulgaris*, S *L. v. vulgaris, L. v. meridionalis*) also have the largest N_e_’s (Fig. 4B, Table S9). This suggests that post-divergence gene flow contributed to elevated N_e_ estimates, although long-term population structure within lineages may also inflate N_e_. We postulate that the existence of a transitional phase, in which differentiated lineages exchange genes, may explain or contribute to the large ancestral population sizes inferred by various methods (IM, ABC, PSMC) in many species (e.g. Duvaux et al., 2011; Nadachowska-Brzyska et al., 2015; Won et al., 2005).

## 4.4 Introgression explains mtDNA paraphyly

With a partially resolved phylogeny, we were able to test statistically for mtDNA introgression suggested previously (Babik et al., 2005; Nadachowska and Babik, 2009; Pabijan et al., 2015). We confirmed introgression in two of the most striking cases (from N *L. v. vulgaris* into *L. montandoni* and S *L. v. vulgaris* into *L. graecus*), effectively explaining the discordance between mtDNA, nuclear genes and morphology, and attributing it to extensive recent and ancient mtDNA introgression. While there is evidence that it has been accompanied by introgression of nuclear genes in the case of *L. montandoni*, we do not have strong evidence of nuclear introgression into *L. graecus*. Better power to reconstruct the extent and timing of nuclear introgression would be obtained with a larger amount of data, particularly from longer genomic regions to explore information contained in haplotype spectra (Harris and Nielsen, 2013).

## 4.5 Taxonomic implications

We note that the application of a rigid taxonomy in which we appropriate scientific names to all nine genetically distinct lineages delimited herein would not fully capture the intricacies of the evolutionary history of *Lissotriton* newts. As stated previously (Wiens, 2007), the delimitation of species poses the problem of thresholding the continuous processes of speciation and lineage amalgamation. Although a number of authors have recognized some of the morphological *L. vulgaris* subspecies as full species (e.g. Dubois and Raffaëlli, 2009; Frost, 2016; Skorinov et al., 2014; Wielstra et al., 2015), these relegations were not based on new data but on inconclusive mtDNA variation, geographic range limits, and morphology.

We delimited four distinct southern taxa for which our data strongly suggest species status: *L. v. graecus* as *L. graecus* (Wolterstorff, 1906), *L. v. kosswigi* as *L. kosswigi* (Freytag, 1955), *L. v. lantzi* as *L. lantzi* (Wolterstorff, 1914) and *L. v. schmidtleri* as *L. schmidtleri* (Raxworthy, 1988) *L. v.* [and not *L. schmidtlerorum*, see Dubois (2007) and Dubois and Raffaëlli (2009)]. We base this assertion on the concordant signals of divergence and independence from allele frequency- and genealogy-based delimitation methods, genealogical sorting, a nearly complete lack of hybrids in well-sampled areas and little evidence for genetic exchange in the recent past. Moreover, *L. kosswigi* and *L. lantzi* are allopatrically distributed (Skorinov et al., 2014; Wielstra et al., 2015), making gene exchange with other lineages improbable in the near future. More extensive sampling in Bulgaria and eastern Greece is needed to characterize the contact between L. schmidtleri and other smooth newt lineages. On the other hand, four central European *L. vulgaris* lineages are interconnected by nontrivial amounts of gene flow. These taxa show fusion following range expansion in the morphologically cryptic N *L. v. vulgaris* and S *L. v. vulgaris*, extensive genetic exchange with neighboring lineages in *L. v. ampelensis*, and continuous gene flow since divergence between *L. v. meridionalis* and S *L. v. vulgaris*. These results suggest that we may be observing the loss of incipient divergence through hybridization in these taxa.

## 4.6 Conclusions

We found that *Lissotriton* newts represent a continuum of genetic divergence with varying levels of shared genetic variation among taxon pairs. This pattern invokes long-term isolation and independent evolution in four southern smooth newt species, genetic exchange across parapatric borders for several taxa in Central Europe, and effective fusion of previously separated gene pools of two lineages in the post-glacial expansion areas of Western Europe. The profound discordance between taxonomic boundaries and mtDNA in *Lissotriton* is explained by extensive mtDNA introgression rather than by other processes. These features make the *L. vulgaris* species group a suitable system to examine the fission/fusion dynamics of the speciation continuum. The newt system is relevant because many species may have experienced such long-term structured populations. However, we also highlight that better methods, jointly modeling incomplete lineage sorting and gene flow, are needed to analyze such systems.

## Acknowledgements

We thank Marcin Piwczyński and Ben Wielstra for insightful comments on the manuscript. Piotr Grodek, Marcin Liana and Krystyna Nadachowska-Brzyska offered invaluable help in the field. Pim Arntzen, Konstantinos Sotiropoulos and Ben Wielstra generously shared samples and locality information. The work was funded by the Polish National Science Centre (2012/04/A/NZ8/00662 to W.B) and the Jagiellonian University (DS/WBiNoZ/INoS/762/15).

## Figure Legends

**Supplementary Figure 1 | Experimental timelines for mouse model pretreatments and *C. difficile* infection.** 9 wild-type C57BL/6 mice across 3 cages were included in each treatment group. **(A)** Streptomycin or **(B)** cefoperazone administered *ad libitum* in drinking water for 5 days with 2 days recovery with untreated drinking water before infection, **(C)** a single clindamycin intraperitoneal injection one day prior to infection, or **(D)** no antibiotic pretreatment (for both SPF control and GF mice). If no antibiotics were administered in the drinking water, mice were given untreated drinking water for the duration of the experiment beginning 7 days prior to infection. At the time of infection, mice were challenged with 1×10^3^ *C. difficile* str. 630 spores. Euthanization and necropsy was done 18 hours post-challenge and cecal content was then collected.

**Supplementary Figure 2 | Analysis of bacterial community structure resulting from antibiotic treatment.** Results from 16S rRNA gene amplicon sequencing from bacterial communities of cecal content in both mock-infected and *C. difficile* 630-infected animals 18 hours post-infection across pretreatment models. **(A)** Non-metric multidimensional scaling (NMDS) ordination based on Theta_YC_ distances for the gut microbiome of all SPF mice used in these experiments (n = 36). All treatment groups are significantly different from each other groups by AMOVA (*P* < 0.001). **(B)** Inverse Simpson diversity for each cecal community from the mice in (A). Cecal communities from mice not treated with any antibiotics are significantly more diverse than any antibiotic-pretreated condition (*P* < 0.001). **(C)** Representation of 16S amplicon reads contributed by *C. difficile* in each sequenced condition compared to the total bacterial community. The percents listed at the top of each group is the proportion of the total community represented by *C. difficile*. Significantly less were for *C. difficile* were detected in each condition (*P* < 0.001).

**Supplementary Figure 3 | Levels of within-group variation across datasets generated for this study. (A)** Normalized transcript abundance of select housekeeping and central metabolism genes. (I) Housekeeping genes; DNA gyrase subunit A (GyrA), threonyl-tRNA synthetase (ThrS), and ATP-dependent Clp protease (ClpP).(II) Genes in separate metabolic pathways that contribute to input substrate score; enolase, glycine reductase (GrdA), and D-proline reductase (PrdA). **(B)** Median sample variance for vegetative *C. difficile* cfu from each colonized condition. **(C)** Median and interquartile range of the sample variance of OTU abundances from 16S rRNA gene sequencing, sample variances for each OTU were calculated individually prior to summary statistic calculations. **(D)** Median and interquartile range of the sample variance of Scaled intensities from untargeted metabolomic analysis, sample variances for each metabolite were in the same fashion as with OTU abundances. Data (other than transcriptomic results) was collected from the same nine animals per group were (n = 9).

**Supplementary Figure 4 | Select *C. difficile* gene set expression compared between treatment group.** Relative abundances of *C. difficile* transcript for specific genes of interest. **(A)** Transcription for select genes from the *C. difficile* sporulation pathway with the greatest variation in expression between the conditions tested. **(B)** Relative abundances of transcript for genes that encode effector proteins from the *C. difficile* pathogenicity locus. **(C)** Transcript abundances for genes associated with quorum sensing in *C. difficile*. **(D)** Transcript relative abundance of select sigma factors which expression or activity is influenced by environmental metabolite concentrations. Asterisks (*) indicate genes from which transcript was undetectable.

**Supplementary Figure 5 | Change in *in vivo* concentrations of additional Stickland fermentation substrates.** Comparison of concentrations for other Stickland fermentation substrates from *C. difficile*-infected and mock-infected mouse cecal content 18 hours post-infection. Labels in the top left corner of each panel indicate whether the amino acid is a Stickland donor or acceptor. Black asterisks inside the panels denote significant differences between mock and *C. difficile*-infected groups within separate treatment groups (all *P* < 0.05). Gray asterisks along the top margin of each panel indicate significant difference from untreated SPF mice (all *P* < 0.05).

**Supplementary Table 1 | Specific genes and normalized cDNA read abundances included in analysis reported in Figure 2.** Transcript abundances reported in each of the antibiotic associated columns were first normalized to both sequencing read length and target gene length. Each of the three groups were then even subsampled to an equal total sequences abundance of 13,000 reads to allow for comparability between groups. Additional columns indicate specific gene annotation (gene, pathways, & KEDD_ID) as well as which group each gene belongs for ternary plot (family).

**Supplementary Table 2 | Normalized cDNA read abundances, gene annotations, and enzymatic reaction information used for metabolic model building for *C. difficile* str. 630 KEGG orthologs across colonized conditions.** All KEGG orthologs included in the *C. difficile* str. 630 KEGG genome annotation (2015) were included in this analysis. Read abundances were normalized as previously outlined to sequencing read length, target gene length, and even total sampling between groups. Also included are individual enzyme annotation for each KEGG ortholog, as well as the associated biochemical reaction information extracted from reaction/reaction_mapformula.lst from KEGG. Together, KEGG ortholog and enzymatic reaction data were used to reconstruct the metabolic network of *C. difficile* str. 630 in presented analyses.

**Supplementary Table 3 | Topology metrics for enzyme and metabolite nodes in the *C. difficile* str. 630 metabolic network.** Topology analysis of the metabolic network assembled for this study was performed in the absence of transcriptomic data to assess quality of *de novo* assembled network in its reflection of known bacterial metabolism patterns. Enzyme and metabolite node analysis are presented on separate tabs. Centrality metrics and brief explanations are as follows: Degree is the total number of connections for a given node (both incoming and outgoing), Betweenness is the number of shortest paths connecting all other nodes pairs that pass through the node of interest, and Closeness is the inverse sum of shortest path length that pass through the node of interest. Combined these calculation inform how strongly connected a node is and how vital it is too overall network structure.

**Supplementary Table 4 | Metabolites with significant metabolite scores for *C. difficile* in each colonized condition.** Each tab represents those metabolites found to exceed the significance cutoffs for *C. difficile* str. 630 after colonization of each of the respective susceptible states. These threshold were set for each metabolite independently through Monte Carlo simulation as outlined by Figure 3C. A *p*-value of < 0.05 corresponded to a metabolite scoring outside of the 95% confidence interval in the random distribution, and *p* < 0.01 corresponds to those outside the 99% confidence interval. Confidence interval calculations for non-normal distributions were performed as defined by (59).

**Supplementary Table 5 | *In vitro* growth analysis for *C. difficile* 630 with carbon sources identified by metabolic network algorithm.** Analysis of growth on highly scored carbon sources to identify possible differences in utilization efficiency.

